# Expanding the genetic toolkit of *Tribolium castaneum*

**DOI:** 10.1101/269464

**Authors:** Johnathan C. Rylee, Dylan J. Siniard, Kaitlin Doucette, Gabriel E. Zentner, Andrew C. Zelhof

## Abstract

The red flour beetle, *Tribolium castaneum*, is an important model insect and agricultural pest.However, many standard genetic tools are lacking or underdeveloped in this system. Here, we present a set of new reagents to augment existing *Tribolium* genetic tools. We demonstrate a new GAL4 driver line that employs the promoter of a ribosomal protein gene to drive expression of a UAS responder in the fat body. We also present a novel dual fluorescent reporter that labels cell membranes and nuclei with different fluorophores for the analysis of cellular morphology. This approach also demonstrates the functionality of the viral T2A peptide for bicistronic gene expression in *Tribolium*. To facilitate classical genetic analysis, we created lines with visible genetic markers by CRISPR-mediated disruption of the *yellow* and *ebony* body color loci with a cassette carrying an attP site, enabling future φC31-mediated integration. Together,the reagents presented here will facilitate more robust genetic analysis in *Tribolium* and serve as a blueprint for the further development of this powerful model’s genetic toolkit.

## Introduction

The red flour beetle (*Tribolium castaneum*), a pest of stored agricultural products, has emerged as a promising system for biological research. It is a representative of the order Coleoptera, which comprises approximately 40% of known insect species and 25% of all known animals [1]. While *Drosophila melanogaster* is by far the most popular insect model system, many aspects of its development and physiology are not representative of insects in general, and so findings in *Tribolium* may be more broadly applicable to insects in many cases. Furthermore, Coleoptera includes significant agricultural pests such as the corn rootworm, Colorado potato beetle, and Asian longhorn beetle, and so using *Tribolium* as an insect model may lead to advances in pest control.

Advances in genetic tools have cemented the status of *Tribolium* as the second model insect of choice behind *Drosophila*. Transgenic *Tribolium* may be obtained using various transposons [2, 3] and more recently via CRISPR/Cas9-mediated genome editing [4]. Eye-specific fluorescent markers have also been developed to aid identification of transgenics [2]. Transposition techniques have been used in for insertional mutagenesis, allowing the identification of essential genes as well as enhancer traps [5]. Another valuable tool for functional genomics, RNA interference (RNAi) via injection of eggs, larvae, or adults, has been implemented in *Tribolium*, both in targeted studies [6-8] and in a large-scale screen of the protein-coding genome [9]. Lastly, the GAL4/UAS system, a popular choice for spatiotemporally controlled expression of a gene of interest in *Drosophila*, has been demonstrated to function in *Tribolium* in the presence of a species-specific basal promoter [10].

Despite the proliferation of tools for genetic analysis and manipulation of *Tribolium*, notable gaps remain. In the case of the GAL4/UAS system, only two driver lines are available, one using a heat shock promoter [10] and the other making use of the odorant receptor co-receptor (*Orco*) regulatory regions [11]. Furthermore, *Tribolium* strains with visible phenotypic markers of known genetic location, which are staples of classical genetic analysis, are not readily available. Here, we present a set of reagents to address these issues and enhance the utility of *Tribolium* as a genetic model organism. We first present a GAL4 driver line that employs a ribosomal gene promoter to direct expression in the fat body and can serve as an second effective marker for transgenesis. In addition, we describe a GAL4-inducible cellular reporter in which the nucleus and endomembrane system are labeled with different fluorescent proteins, acting as a robust means by which to analyze cellular structure, particularly with respect to neurons. Furthermore, both the GAL4 and UAS cloning vectors are designed to accept any gene or genomic region of interest to generate new drivers and reporters. We also address the lack of visible phenotypic markers in *Tribolium* by using CRISPR to disrupt two genes involved in cuticle pigmentation via homologous recombination with cassettes containing an attP site to facilitate future genomic insertion of DNA of interest using the φC31 integrase. The tools presented here represent a valuable resource for the *Tribolium* research community and serve as a general template upon which further tools can be based.

## Materials and methods

### Tribolium husbandry and strains

All animals were raised at 28^°^C on a standard flour yeast mix. The following strains were utilized: *v*^*w*^ and *m26*[3].

### Vectors

All vectors will be made available through the *Drosophila* Genomics Resource Center at Indiana Universityhttp://dgrc.bio.indiana.edu/Home).

#### P119der

119der, a derivative of pSLfa[UAS-Tc’Hsp-p-tGFP-SV40] (kindly provided by Dr. Gregor Bucher)[10], was constructed by excision of tGFP with KpnI and NotI followed by replacement with the sequence ACTAGTGAATTCAAAGTACCACTCGAGAGCGGCCGCG. This replacement destroyed the KpnI site but preserved the unique NotI site and added a unique XhoI site.(DGRC # XXXX)

#### P130der

p130der, a derivative of pSLfa[Hsp-p-Gal4Delta-SV40_attp] (kindly provided by Dr. Gregor Bucher) [10], was constructed by digestion with BamHI and addition of the sequence GGATCCAGGTACCAGCGGCCGCAGGATCC, containing unique KpnI and NotI sites. (DGRC # XXXX)

#### pGZ286 (pBac-3xP3-EGFP-pTC006550-GAL4Δ)

The basal *hsp68* promoter of PCR-linearized p130der was replaced with the *TC006550* promoter amplified from blaAmp-Tc6550Pro-GFPZeo-Luciferase-HSP-Orange-pIZT (kindly provided by Dr. Yoonseong Park) [12] with NEBuilder HiFi assembly (NEB). The resulting p*TC006550*-*GAL4Δ*-SV40 polyA coding sequence was then amplified and inserted into PCR-linearized pBac[3xP3-EGFP] with NEBuilder HiFi assembly. (DGRC # XXXX)

#### pTC241 (pBac-3XP3-EGFP-UAS-nls-EGFP-T2A-mCherry)

The nls-EGFP-T2A-mCherryCAAX insert was amplified from pSYC-102 (a gift from Seok-Yong Choi, Addgene plasmid #74790) [13] as two fragments, and assembled into the NotI site of p119der using NEBuilder HiFi assembly. UAS-nls-EGFP-T2A-mCherryCAAX was then excised using flanking AscI sites and ligated into pBac[3xP3-GFP]. (DGRC # XXXX)105

### Immunofluorescence and imaging

*Tribolium* larvae were dissected in PBS and tissue was fixed in 4% paraformaldehyde/PBS. The tissue was washed 2X with PBT (PBS and 0.8% Triton-X 100) and incubated with DAPI (final concentration of 0.1 µg/ml) for 10 minutes and then washed with PBS before mounting for imaging (modified from [14]). Confocal images were captured on a Leica SP8 confocal utilizing a Leica 63X of oil immersion objective with a numerical aperture of 1.40. Light and fluorescent whole animal images were collected on a LeicaMZ10F dissecting microscope. All pictures were processed in Adobe Photoshop.

### CRISPR

115 gRNAs were designed using CRISPRdirect [15] using the Tcas3 genome assembly for the specificity check. Only high-quality gRNAs were selected. 20-mer protospacer sequences were cloned into BsaI-digested pU6b-BsaI-gRNA [4] by NEBuilder HiFi-mediated ssDNA oligo bridging as desbribed (http://www.neb.com/-/media/nebus/files/application-notes/construction-of-an-sgrna-cas9-expression-vector-via-single-stranded-dna-oligo-bridging-of-double-stranded-dna-fragments.pdf) using an ssDNA oligo consisting of the protospacer flanked by 25 bp regions of vector homology. Protospacer sequences used were CCGGAAAATAATCTCCCAGT (*yellow*, *TC000802*) and TTTCGTAAAAGTTTGAATCG (*ebony*, *TC0011976*) (DGRC #XXXX and#XXXX). Homology donors consisting of an attP site and 3xP3-DsRed-SV40 polyA flanked by loxP sites in the same orientation between 800 bp (*yellow*; left arm: ChLG2:7,663,877-7,664,676, right arm: ChLG2:7,664,677-7,665,476) or 726/739 bp (*ebony*; left arm:ChLG9:13,340,543-13,341,268, right arm: ChLG9:13,341,269-13,342,007) homology arms were synthesized by IDT and delivered in pUCIDT-amp. Mixtures consisting of 400-500 ng/mL each of Cas9 plasmid [4], sgRNA plasmid, and donor plasmid were injected into *v*^*w*^ embryos.

## Results

### Construction and implementation of a *ribo*-GAL4 driver

Inducible expression of a transgene of interest is a key capability in any genetic model organism. In *Drosophila*, the bipartite *Saccharomyces cerevisiae* GAL4/UAS (upstream activating sequence) system, in which the GAL4 transcription factor binds to a 17-bp motif within the UAS to drive transcriptional activation, is frequently used for spatiotemporally controlled expression of a gene of interest [16]. In *Tribolium*, the GAL4-UAS system has been demonstrated to be functional when the UAS is coupled with the *hsp68* basal promoter and driven by heat shock-inducible GAL4 [10]. However, no ubiquitous GAL4 drivers have been reported for *Tribolium*. Such drivers are useful when screening for an organismal or developmental phenotype of overexpression or knockdown.

In an attempt to generate a ubiquitous and constitutive GAL4 driver line, we considered a previous study in which the promoter of a ribosomal protein gene (*TC006550*) was used to drive high-level expression in the *Tribolium* TcA cell line [12]. We thus replaced the heat shock promoter of our p130der vector (see Materials and Methods), bearing the basal *hsp68* promoter and GAL4Δ, with the *TC006550* promoter. GAL4Δ is a variant of GAL4 in which the N- and C-termini, containing the DNA-binding and transcriptional activation domains, are directly fused [17]. This variant of GAL4 has been shown to increase transactivation by ~2-fold in *Drosophila*[18]. We then transferred the *TC006550* promoter and GAL4Δ coding region to a piggyBac vector containing 3xP3-EGFP, enabling selection of transgenics by fluorescence in photoreceptors. This vector was injected into a *Tribolium* line lacking eye pigmentation (*vermillion^white^* (*v*^*w*^)) with 3xP3-DsRed-marked piggyBac transposase integrated into the X chromosome [3]. Resulting adults were outcrossed to *v*^*w*^ and progeny were assessed for GFP expression in the retina. *TC006550-GAL4Δ* transformants (hereafter referred to as *ribo*-GAL4) were identified and the DsRed-marked transposase was removed through subsequent crosses and two independent insertions were generated.

To assay the functionality of the *ribo*-GAL4 driver line, we crossed it to a previously described UAS-GFP responder line [10]. Larvae displayed strong whole-body fluorescence (Fig 1A-C) and fluorescence is maintained throughout pupal development and into adulthood as expected for a ubiquitous expression. However, further examination revealed fluorescence was only detected in the putative fat body (Fig 1D-G) and absent from other tissues (e.g. the gut, muscle, and CNS). We speculate that the lack of ubiquitous GAL4 expression in the *ribo*-GAL4 line may reflect tissue-specific differences in ribosomal protein gene expression [19] and that, given the apparent *in vivo* expression profile of *TC006550*, the TcA cell line may be derived from fat cells.

**Fig. 1.**
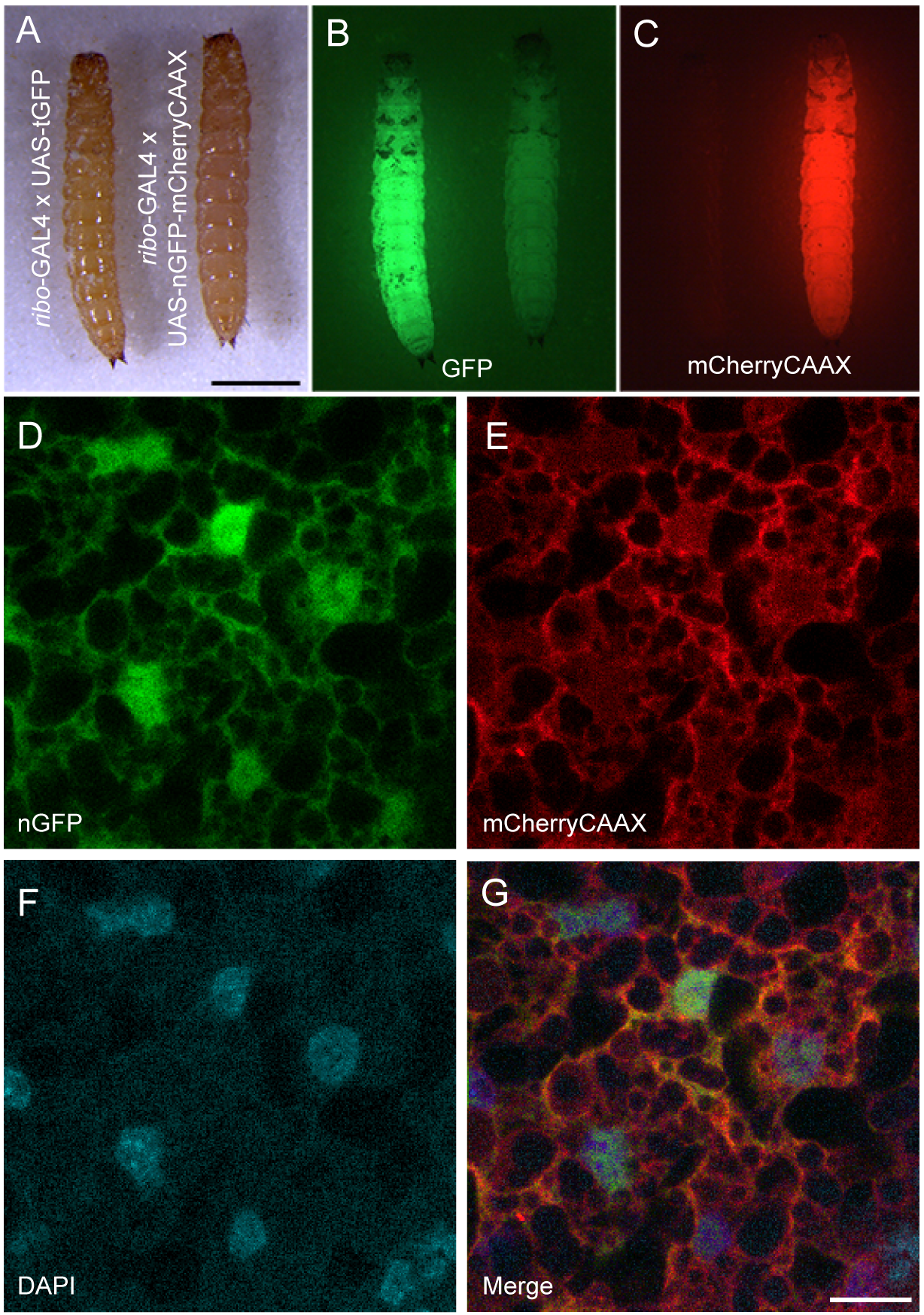
Characterization of GAL4 and UAS vectors. (A-C) Tribolium larvae expressing either GFP (left) or nls-EGFP-T2A-mCherryCAAX (right) driven by *ribo*-GAL4. (A) White light illumination; (B) GFP illumination. The nls-EGFP signal is not detectable as compared to cytoplasmic GFP. (C) mCherry illumination. Scale bar = 1 mm. The mCherry illumination mimics the cytoplasmic GFP expression. (D-G) Images of fat cells expressing nls-EGFP (D) and mCherryCAAX (E), counterstained with DAPI (F), and the merge of the three labels (G). Each represents a single confocal section. Scale bar = 10 μm.

### Construction and implementation of a reporter for cell structure

There are numerous reporters available for highlighting cell structure and function. Our goal was to test whether these reporters could be simply swapped into a universal UAS cloning vector for *Tribolium* with zero or minimal changes to the already existing sequence. We chose nls-EGFP-T2A-mCherryCAAX [13]to test the utility of bicistronic fluorescent reporter expression for studying cell morphology as well as the use of the viral T2A peptide in *Tribolium*. When combined with our *ribo*-GAL4 line, mCherry expression could easily be detected in whole larvae, mimicking the spatial and temporal pattern obtained with cytoplasmic GFP (Fig 1A-C). To confirm the expression and localization of both the nuclear GFP and the endomembrane linked mCherry we examined the subcellular localization of each in fat cells. Colocalization with DAPI confirmed the subcellular localization of GFP in the nucleus with mCherryCAAX bound to membranes (Fig 1D-G). These results indicate that existing fluorescent reporters can be easily implemented in *Tribolium* using our UAS vector and that the T2A peptide can be used for multicistronic gene expression in *Tribolium*.

### CRISPR-based generation of lines with visible phenotypic markers

Defined phenotypic visible markers are essential for facilitating even basic tasks such as establishment of stocks of genetically modified organisms or tracking of specific chromosomes through multiple generations. While the scope of genetic reagents available in *Tribolium* has markedly increased over the past few decades, there are notable limitations and gaps in existing tools. For instance, the use of visible phenotypic markers for genetic mapping is quite limited, especially compared to *Drosophila*, where variations in wings, bristles, eye and body pigmentation, and body shape are available. Additionally, while dominant mutations giving rise to visible phenotypes in *Tribolium* have been documented [20-22], they have not been mapped
to a tractable genetic interval in most cases and so cannot be used in such analyses.

In order to create a general strategy for the expansion of a pool of visible phenotypic markers for *Tribolium*, we employed CRISPR based homologous recombination. We designed gRNAs against the coding regions of the *yellow* (*TC000802*) and *ebony* (*TC011976*) genes, as well as a disruption cassette with useful genetic features. In many insects, the disruption of either *ebony* and *yellow* leads to viable and fertile animals that are easily identifiable from their wild-type counterparts. With respect to our CRISPR procedure, we chose a homologous recombination strategy that would permit detection of a disruption in either locus regardless of whether a change in pigmentation resulted. For each targeting construct, we utilized 700-800bp homology arms, and between them we enclosed an attP site to facilitate future φC31 integrase-mediated insertion of DNA of interest at a defined location [23]. More importantly, we included the DsRed Express fluorescent protein under the control of the eye-specific 3xP3 regulatory element to facilitate screening of CRISPR based recombinants; and two loxP sites in the same orientation, enabling future Cre-mediated excision of 3xP3-DsRed from the genome (Fig. 2A). The positioning of the loxP sites is such that they flank only the fluorescent marker and so excision would leave the attP site intact, effectively recycling the DsRed marker for further use.

**Fig. 2.**
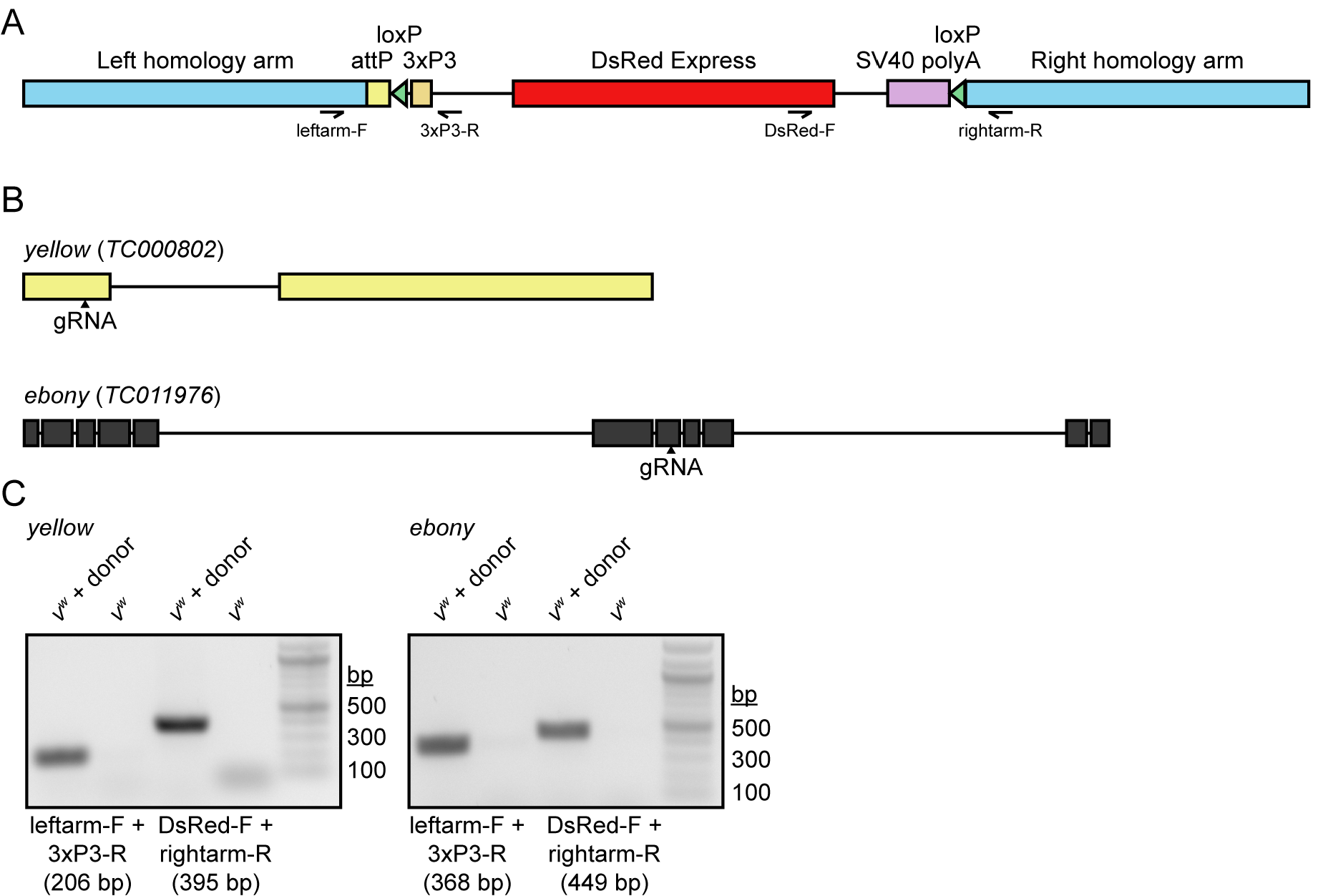
Construction and validation of the CRISPR gene disruption strategy. (A) Schematic representation of the gene disruption cassette used in this study. The positions and directionalities of primers used for genomic PCR validation are included in the schematic. (B) Positions of gRNAs used to disrupt the *yellow* and *ebony* loci. (C) Genomic PCR demonstrating the presence of each gene disruption cassette in the genome of transgenic but not parental beetles.

*v*^*w*^ embryos were injected with a mixture of Cas9 plasmids, gRNA expression plasmid, and repair template plasmids. Surviving adults were crossed to *v*^*w*^ and resulting progeny were screened upon eclosion for DsRed expression in the retina. For *yellow*, 308/356 injected larvae survived to adulthood and of these, two tested positive for germline transmission of the disrupted gene (0.65%). For *ebony*, 184/217 injected larvae survived to adulthood and of these, one tested positive for germline transmission of the disrupted gene (0.54%).

Characterization of *yellow*-edited adults revealed that cuticles of newly eclosed beetles were noticeably lighter than those of their *v*^*w*^ counterparts, but their color darkened over time until they were difficult to distinguish from the parental line (Fig 3). In contrast, adults with homozygous disruption of *ebony* displayed substantially darker cuticular pigmentation than parental *v*^*w*^ individuals (Fig. 4A-B) and DsRed fluorescence in their eyes (Fig. 4). While *ebony* CRISPR individuals were clearly phenotypically distinct from the parental line, their cuticle was not as dark as individuals injected with *ebony* dsRNA (http://ibeetle-base.uni-goettingen.de/details/TC011976). We speculate that this difference is due to the location of the gRNA target site. It lies within the seventh exon of *ebony*, which falls after the sequences encoding all but one predicted functional domain of the protein (Fig. 5). However, this gRNA falls at the start of the last domain of the protein, and disruption of this domain likely explains the hypomorphic phenotype observed.

**Fig. 3.**
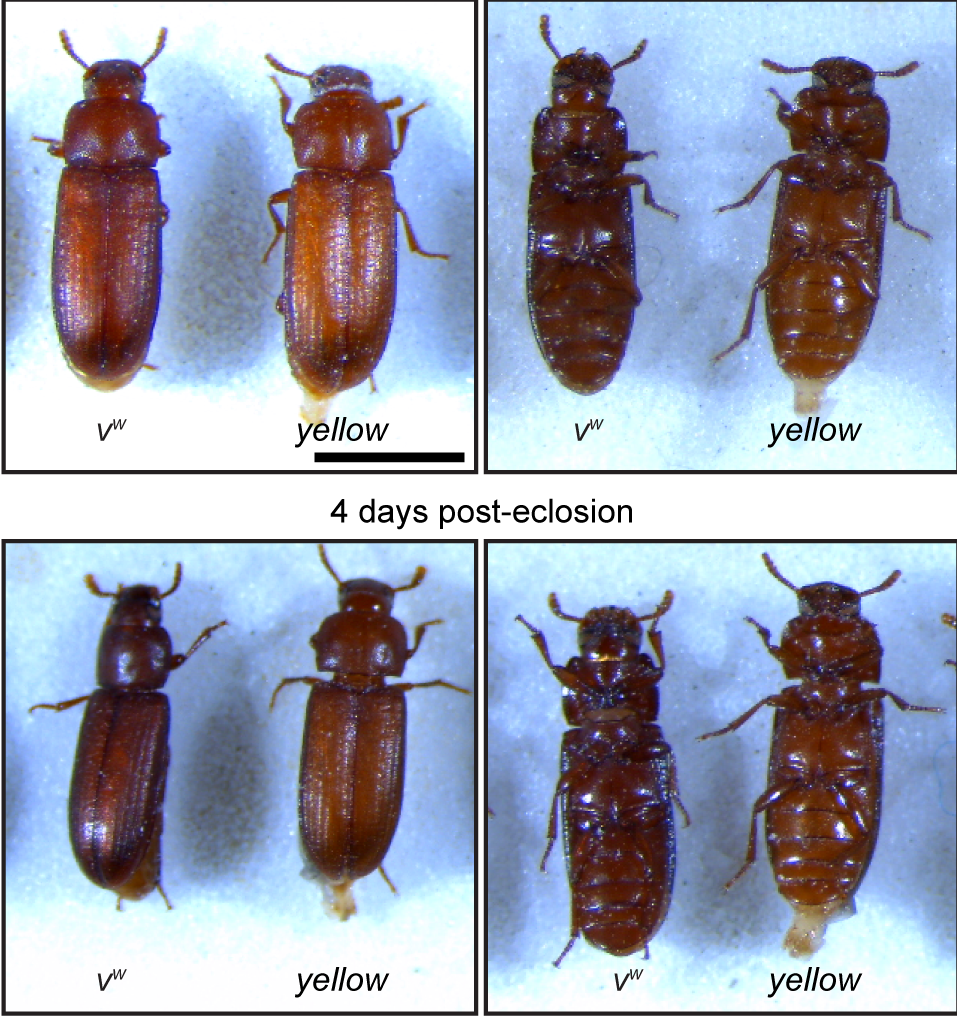
Phenotypic characterization of yellow CRISPR beetles. Dorsal and ventral views of parental *v*^*w*^ and transgenic *yellow* CRISPR beetles at 2 and 4 days post-eclosion. Scale bar = 1 mm.

**Fig. 4.**
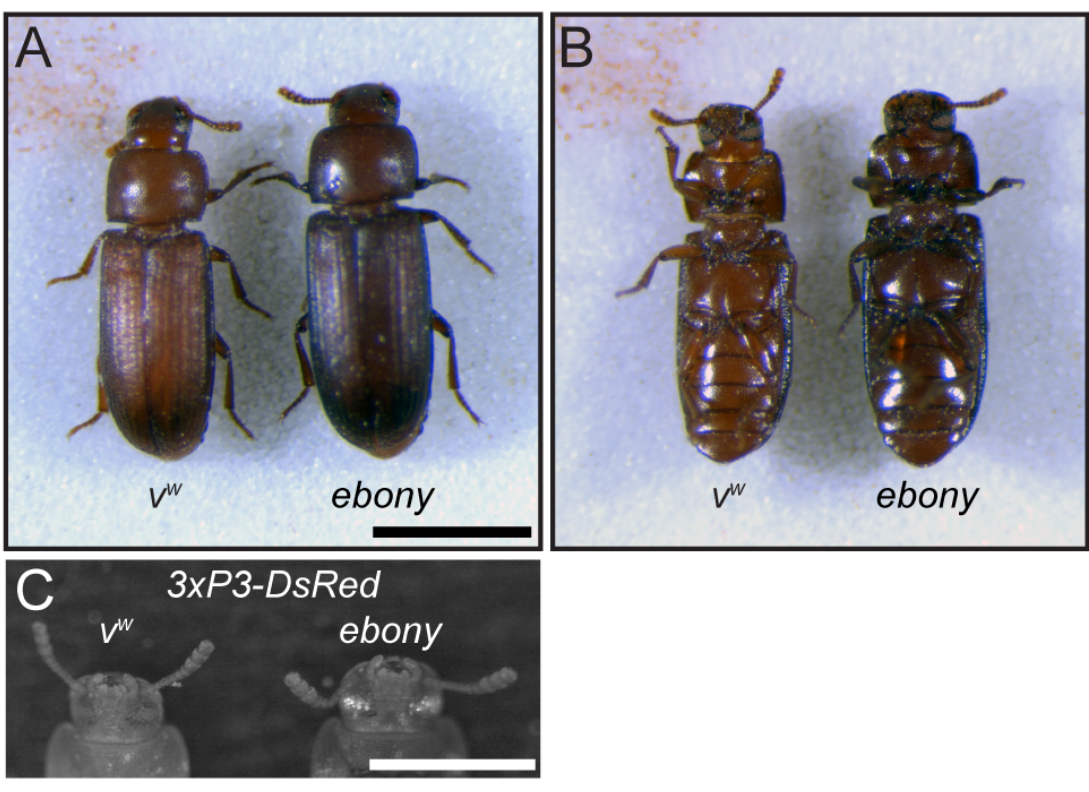
Phenotypic characterization of *ebony* CRISPR beetles. (A) Dorsal and (B) ventral views of parental *v*^*w*^ and transgenic *ebony* CRISPR beetles. Scale bar = 1 mm. (C) DsRed fluorescence microscopic image of the eyes of *v*^*w*^ and transgenic *ebony* CRISPR beetles. Scale bar = 0.5 mm.

**Fig. 5.**
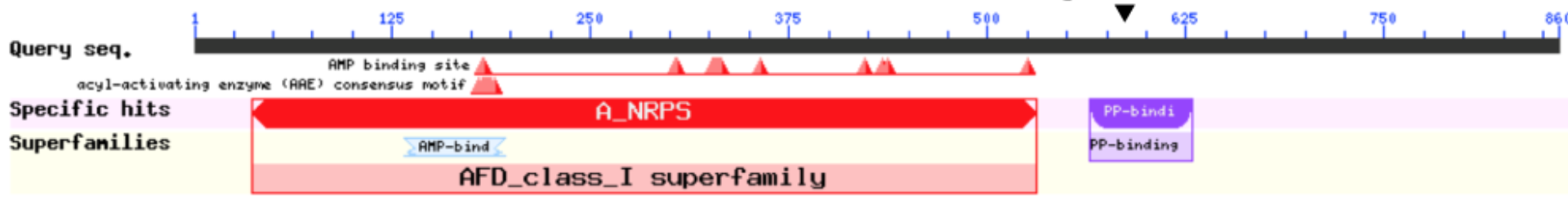
NCBI conserved domain search results for the *ebony* protein. The location of the gRNA relative to the mature protein sequence is indicated.

## Conclusions

Here, we present a series of reagents aimed at increasing the genetic tractability of *Tribolium*. These are (1) a GAL4 driver plasmid and *Tribolium* line expressing GAL4 in fat cells; (2) a UAS plasmid and UAS-inducible dual-nuclear/endomembrane fluorescence reporter for analyzing cell structure; (3) a template gene disruption cassette for the generation of mutants and insertion of attP sites; and (4) *Tribolium* lines carrying disruptions of two body color loci, *yellow* and *ebony*.

We surmise that the failure of the *ribo*-GAL4 line to drive ubiquitous reporter expression is attributable to its late pupal origin [24], reflecting cell type variability in *TC006550* expression. Indeed, heterogeneity in ribosomal protein expression across cell types has been widely reported [19, 25-27]. Other potential candidates for the establishment of a ubiquitous GAL4 line include the *α-Tubulin1* promoter, which has been shown to drive ubiquitous GFP expression throughout the *Tribolium* life cycle [28], and the *Polyubiquitin* promoter [29]. The p130der plasmid permits efficient insertion of any potential genomic sequence for designing future GAL4 lines. Furthermore, our data demonstrates that any established reporter can be cloned into our modified version of pSLfa[UAS-Tc’Hsp-p-tGFP-SV40] [10], p119der, for direct expression in *Tribolium*. Future variations/deviations of both p130der and p119der will include an attB site for direct insertion into known genomic positions as well as a fluorescent marker to enable rapid screening of transgenics.

While we successfully disrupted the *yellow* and *ebony* loci via CRISPR, the visible phenotypes associated with these editing events were unpredictable. In particular, disruption of *yellow* resulted in a slow-tanning phenotype, with young adults displaying a visibly lighter cuticle than the parental strain that then darkened until it was indistinguishable from that of non-edited beetles. In the case of *ebony*, we were able to achieve a marked darkening of the cuticle using our disruption strategy, but our mutation was potentially hypomorphic when compared to *ebony* RNAi, which yielded a darker black-body phenotype. In order to maximise the visible phenotypes obtainable with CRISPR disruption of an eye or body color locus, we therefore recommend pre-screening of candidate visible marker genes with RNAi prior to initiating genome editing. Several candidates for body eye and body color genes have been assessed by RNAi in the literature [30-33] and may serve as suitable targets for our CRISPR disruption approach. Moreover, our data demonstrate that the inclusion of an independent marker for CRISPR-based modifications is invaluable in recovering transformants and thus can mitigate the uncertainty associated with the targeting of other potential candidate loci for visible markers.

## Acknowledgments

We thank Drs. G. Bucher, S. Brown, and M. Lorenzen for *Tribolium* reagents. We thank Dr. J. Powers and the Indiana University Light Microscopy Imaging Center for assistance with image generation. We thank Dr. M. J. Wade for helpful discussions throughout the course of this work. This work was supported by Indiana University startup funds to G.E.Z. and the NSF (IOS-1353267) to A.C.Z.

---

**S1 Table.**
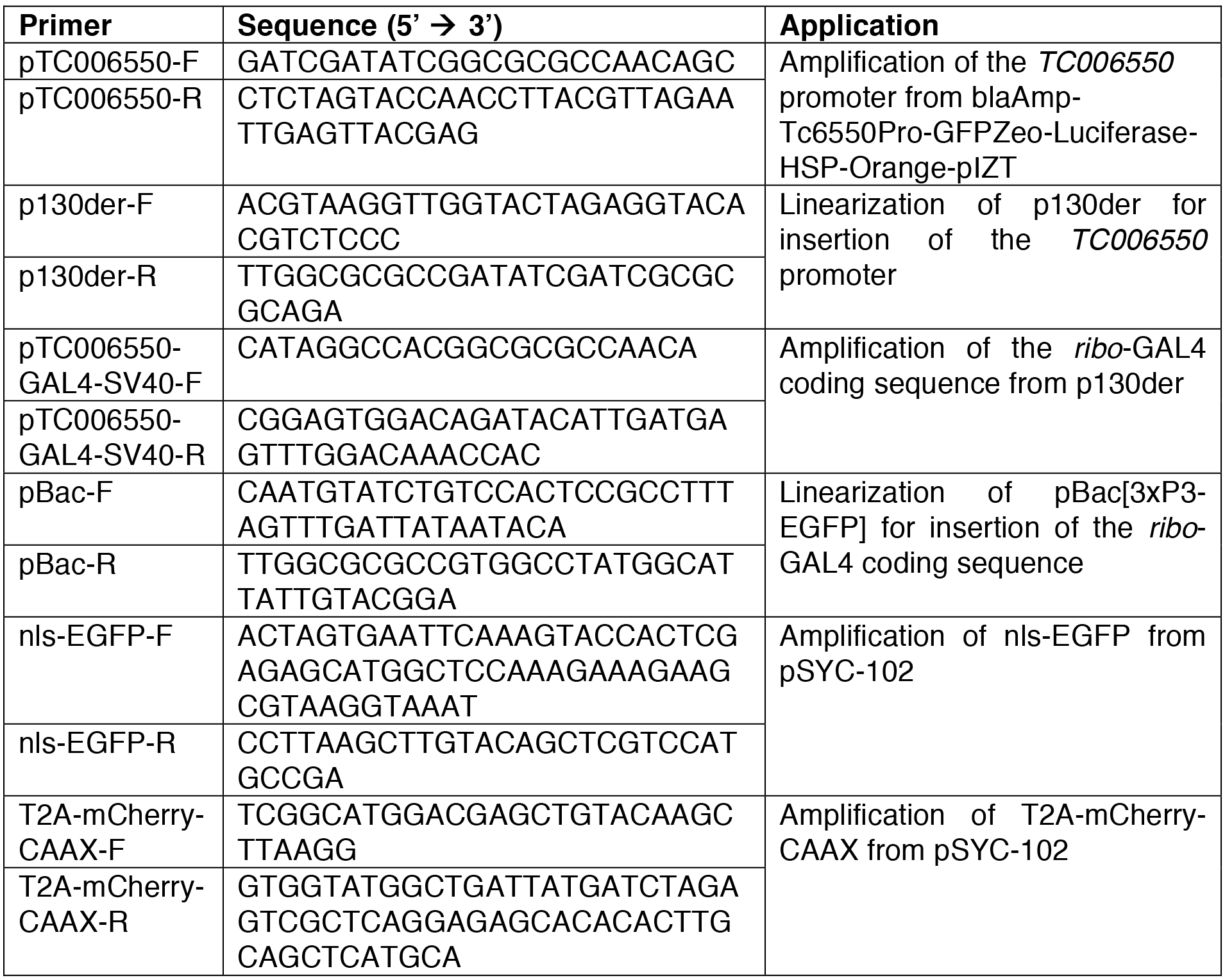
DNA oligonucleotides used in this work.

